# ProteoMapper: Alignment-Aware Identification and Quantitative Analysis of Contextual Motif–Domain Patterns in Protein Families

**DOI:** 10.64898/2026.02.17.706295

**Authors:** Sifullah Mahmud Sefa, Joyeeta Sarkar, Arif Hasan Khan Robin, Machbah Uddin

## Abstract

Protein function depends on interactions between structural domains and regulatory motifs. Yet current tools analyze these elements separately, hindering investigation of disease mutations affecting evolutionarily conserved, structurally constrained motifs. We present ProteoMapper, a computational framework integrating HMMER-based domain annotation with user-defined motif detection to quantify motif-domain spatial relationships in protein families. ProteoMapper introduces two discovery metrics: (1) positional conservation scoring, identifying motifs at identical alignment coordinates in ≥ *N*% of sequences (default 60%), indicating purifying selection; (2) Motif-Domain Coverage Score (MDCS), quantifying motif embedding within Pfam domains (MDCS=1: fully embedded; MDCS=0: extra-domain). The platform processes Excel-formatted alignments without programming requirements, delivering color-coded reports with conserved motif positions, domain boundaries, and MDCS values. Parallel execution of sequence batches enables rapid analysis (8 motifs were searched in 150 sequences with complete Pfam scanning in <6 seconds on standard hardware). Validation across three protein families confirmed technical accuracy and biological insight. In PLATZ transcription factors (24 proteins), domain predictions achieved 0.94 mean intersection-over-union versus published annotations, exactly reproducing 22 of 23 reported spans. In *Arabidopsis* ERD6-like sugar transporters (17 proteins), MDCS analysis revealed canonical PROSITE signatures PS00216 and PS00217 are equally domain-embedded (MDCS=1.0) but evolutionarily divergent. PS00217 shows positional conservation (58.8% of sequences) while PS00216 exhibits dispersal, suggesting subfunctionalization. In tomato actin-depolymerizing factors (11 proteins), domain detection achieved 100% sensitivity with >93% positional concordance. ProteoMapper enables hypothesis-driven investigation of evolutionary constraints, regulatory mechanisms, and variant effect prediction in biomedical and functional proteomics. Source code, documentation, and test results with datasets at https://github.com/sifullah0/ProteoMapper.

## 1 Introduction

Protein function emerges from the coordinated action of conserved structural domains and short linear motifs (SLiMs). Conserved domains provide stable architectural frameworks that enable catalytic activity, molecular recognition, and structural integrity, whereas SLiMs encode compact regulatory logic such as phosphorylation sites, degradation signals, and transient protein–protein interaction interfaces [29, 8, 9]. Together, these elements shape protein behavior across diverse cellular contexts. A central challenge in comparative and functional genomics is determining whether specific sequence motifs are evolutionarily constrained through structural embedding within conserved domains, or whether they evolve independently as flexible regulatory elements. This distinction has important implications for functional interpretation of sequence variation, as motifs that are both positionally conserved and domain-embedded are more likely to contribute directly to core protein mechanisms than motifs located in unconstrained regions [15, 14].

Despite their biological importance, domain and motif analyses are typically performed using separate computational workflows. Profile hidden Markov model (HMM)–based tools such as HMMER and InterProScan robustly identify conserved domains through sequence– profile alignments against curated databases [19, 1], but do not support user-defined motif detection or quantitative assessment of motif–domain spatial relationships. Conversely, motif-focused resources such as ScanProsite, MEME, ELM, and EMBOSS Fuzzpro detect linear sequence patterns but lack integrated domain context, alignment-level conservation analysis, and standardized embedding metrics [7, 3, 8, 26]. Recent methods including MAFin and Wregex advance motif detection through probabilistic scoring and weighted regular expressions [18, 21], yet remain focused on motif identification rather than unified domain-context interpretation. Consequently, researchers must manually integrate outputs from multiple tools, extracting domain coordinates from HMMER `domtblout` files, motif positions from web servers, and inferring overlaps using custom scripts or spreadsheets. This manual integration is error-prone, difficult to reproduce, and provides no quantitative framework for distinguishing motifs that are fully embedded within domain cores from those that overlap domain boundaries or reside in interdomain regions [10].

To address these limitations, we developed *ProteoMapper*, an integrated computational framework for alignment-aware interpretation of protein domains and motifs. The platform introduces two complementary discovery metrics. First, positional conservation scoring identifies motif occurrences fixed at identical alignment coordinates across a user-defined proportion of sequences (default 60%), highlighting motifs under purifying selection for positional constraint. Second, the Motif–Domain Coverage Score (MDCS) quantifies the fraction of each detected motif that overlaps predicted Pfam domains, yielding values between 0 and 1 that distinguish fully embedded, boundary-spanning, and extra-domain motifs. Together, these metrics transform descriptive annotation into quantitative, hypothesis-driven analysis of motif constraint and domain integration.

ProteoMapper operates directly on Excel-formatted multiple-sequence alignments with automatic header detection and integrates HMMER-based Pfam scanning with flexible regular-expression motif detection. The framework produces interpretable, multi-sheet Excel outputs that overlay conserved motif positions, domain boundaries, and MDCS values without requiring programming expertise. We demonstrate the utility of this integrated approach through case studies on PLATZ transcription factors, actin-depolymerizing factors, and ERD6-like sugar transporters, illustrating how combined conservation and domain-context analysis enables systematic investigation of protein architecture, evolutionary constraint, and regulatory organization.

## 2 Experimental Procedures

In this section, various prerequisites are discussed, including dataset collection and preparation, the main algorithms, and the necessary tools for software development. The general process of the proposed model is presented in Fig. 1.

**Figure 1:**
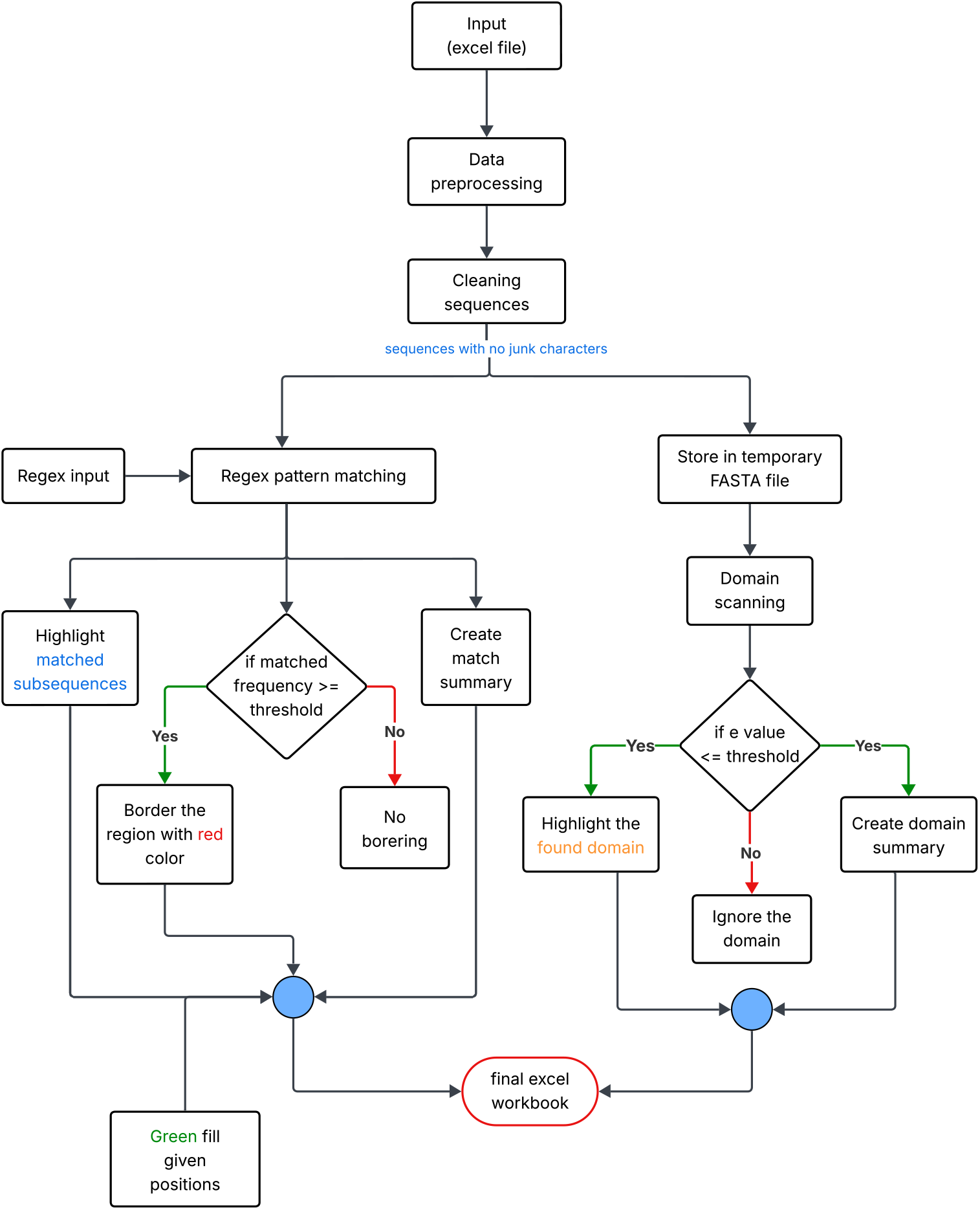
Workflow diagram of the proposed sequence analysis toolkit.

### 2.1 Data collection and preprocessing

ProteoMapper accepts protein sequences in a single Excel file (.xlsx) rather than plaintext formats. This file format offers several advantages for experimental biologists. Excel allows gene names or identifiers to be stored in the same row as their corresponding protein sequences, which prevents synchronization issues that arise when metadata and sequences are maintained in separate files. If sequences include gap characters (for example, “-”), the spreadsheet grid makes alignments easy to view during manual checks. In addition, Excel is widely used in biological research workflows, which reduces the barrier to adoption for experimental users. The input file must contain, at a minimum, two columns: Gene Name / Gene ID, which holds sequence identifiers, and Protein Sequences, which contain protein sequences. To ensure compatibility with diverse user-generated spreadsheets, the header row is detected dynamically by scanning for keywords such as “Protein sequences,” “Gene name,” or “Gene ID.” This automatic detection supports flexible input structures while preserving consistent downstream processing.

#### Sequence cleaning and normalization

Before analysis, raw protein sequences are processed to remove whitespace, line breaks, special characters, or FASTA headers (“>“) that could interfere with motif detection and domain annotation. The following procedures are applied to each input sequence:

- **FASTA handling**. If a sequence begins with “>“, the symbol is removed and a flag records the original header to preserve provenance.
- **Invalid-character removal**. Any character not among the 20 standard amino acids (A, C, D, E, F, G, H, I, K, L, M, N, P, Q, R, S, T, V, W, Y) is deleted, eliminating stray symbols, digits, or other non-amino inputs.
- **Gap-aware normalization**. Two representations are generated:
  – **Display view:** Retains alignment gaps (“-”) for visualization and inspection of alignments.
  – **Matching view:** Removes gaps and any remaining non-amino characters to produce a contiguous sequence suitable for pattern matching and motif searches.
- **Missing-data handling**. Null or missing input values are replaced with empty strings to ensure consistent string operations and avoid downstream errors.

These procedures improve reproducibility and accuracy of subsequent motif detection and domain annotation. Given that multiple-sequence alignments often introduce variable gap patterns, and that algorithms such as MAFFT [11] illustrate how alignment strategies may impact gap distributions and normalization across sequences. By maintaining both a display view (with alignment gaps) and a matching view (without gaps), it is ensured that downstream motif searches and domain annotations are operated on clean, consistent sequences.

### 2.2 Pattern Matching and Highlighting

This toolkit supports flexible detection of protein motifs using user-defined regular expressions. Unlike fixed pattern-matching methods, the regex engine gives researchers full control over the motif patterns they wish to search. Users may define specific or degenerate motifs, such as phosphorylation sites (e.g., `[ST]..[DE]`) or family-specific signatures that are not present in standard databases. Modification sites such as N-glycosylation 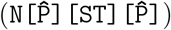 can be identified as well. Patterns are entered directly through the graphical interface (GUI).

The motif-matching engine operates in two modes. For datasets with fewer than 100 sequences, the system uses serial execution. For larger datasets, it distributes computations across multiple worker processes. Users can set the number of processes directly through the interface. Complete algorithmic details are provided in algorithm 1

#### Conservation Threshold

Users can specify the minimum percentage of sequences that must contain a given motif within the same positional boundary, then the motif is classified as conserved within that range. The default threshold is 60%. Motifs that meet or exceed this level are highlighted in the output with a red border.

##### Algorithm 1: Motif Matching and Conservation Highlighting

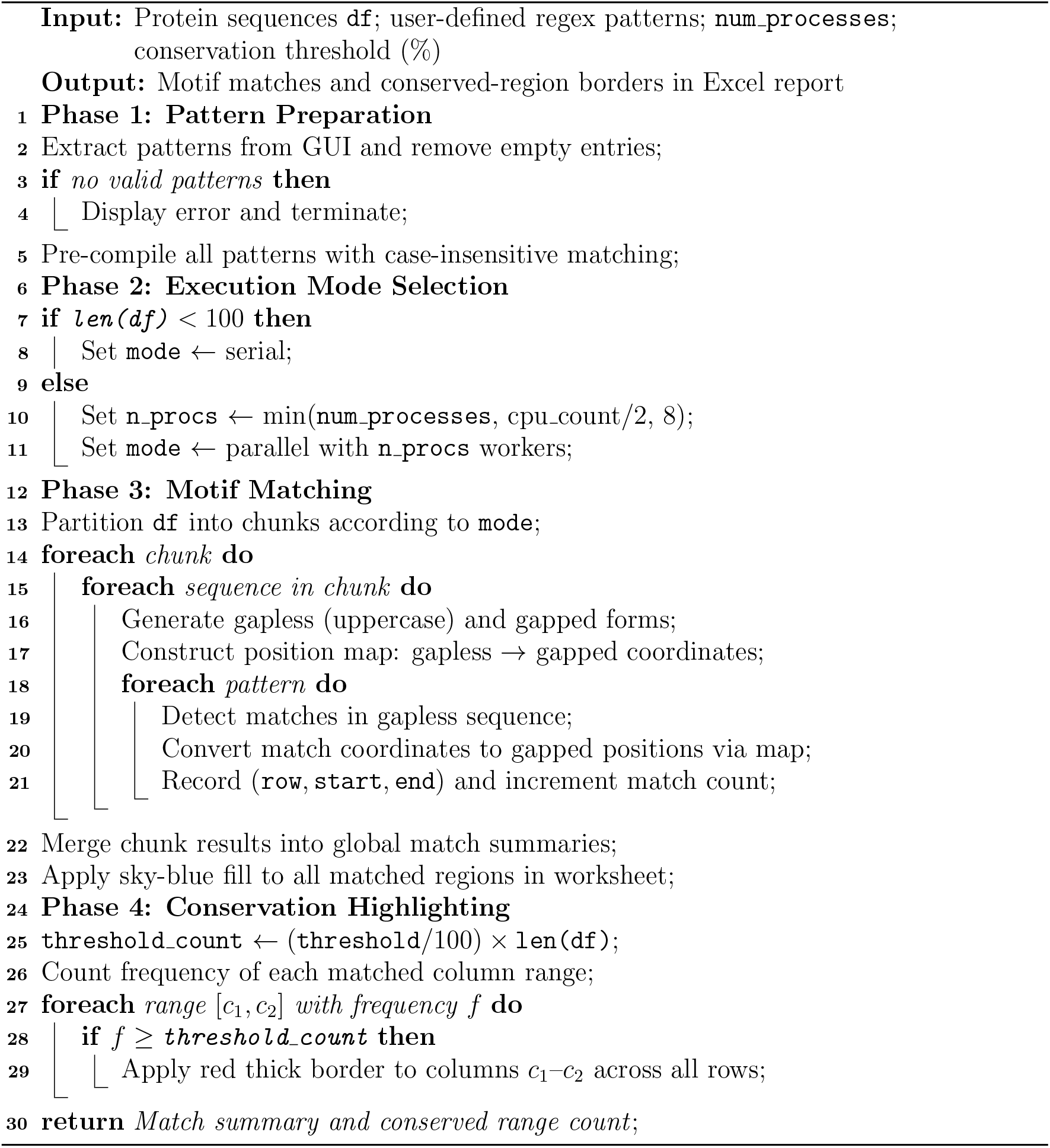

#### Excel Visualization and Error Handling

Outputs are presented in excel workbook with highlighted results. **Sky Blue Fill:** All detected motif occurrences, **Red Border:** Motifs meeting or exceeding the conservation threshold, and **Green Fill:** Specific columns, given by user, listed for targeted review. Similarly, in case of error Handling and edge cases identification, several criteria have been followed. Those are invalid regex patterns are flagged immediately during validation, alignment gaps (-) are automatically ignored in pattern searches, and multiline regex is supported for detecting complex or structural motifs.

### 2.3 Conserved Domain Prediction Using HMMER

The pipeline can process large sequence sets more efficiently through parallel execution. In this mode, multiple `hmmscan` processes run simultaneously. Each process scans a different subset of the input sequences. The results from all processes are then combined into a single output file. First, input sequences are cleaned (gaps removed) and saved to a temporary FASTA file. Each sequence is assigned a numeric ID that maps to its original alignment position for full traceability. Then, `hmmscan` is executed with the `domtblout` output format to perform sequence–profile alignments against Pfam-A. The pipeline performs error checks to detect missing executables or execution failures. Then, Hits are filtered to retain only high-confidence matches (independent E-value ≤ 0.001). Parsed fields include query ID, Pfam accession (e.g., `PF00071`), domain start/end positions, and the bit score. Finally, structured results are written to `hitdata.txt` with consistent delimiters. The pseudocode for all of these processes is presented in algorithm 2. Full domain descriptions are preserved,, and E-values are presented in scientific notation for accuracy. The output is designed for reproducibility and easy tracing back to the original alignment.

#### Algorithm 2: Parallel Domain Detection via Multiple hmmscan Instances

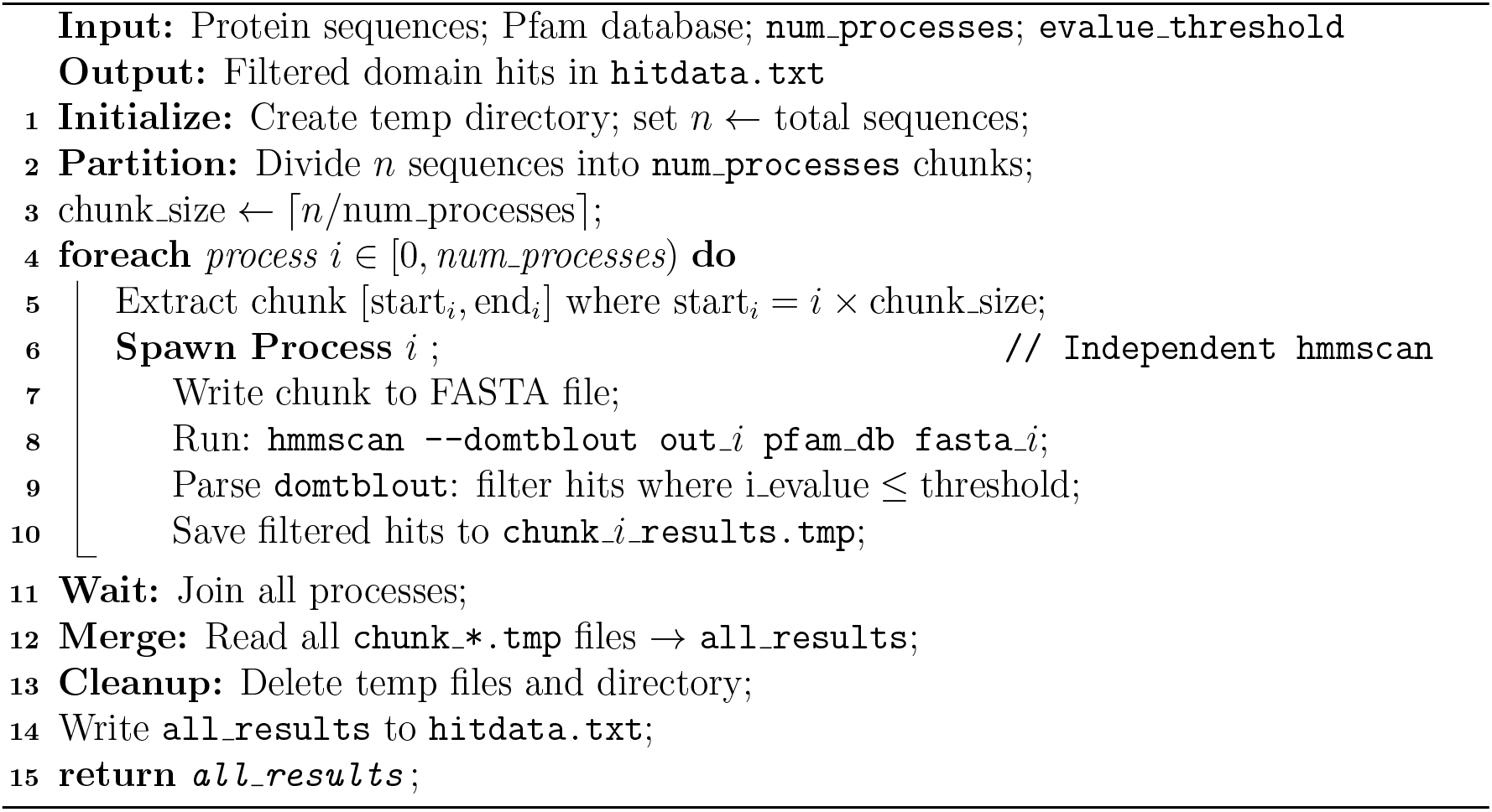

#### Domain Highlighting in Excel

The pipeline generates interactive domain annotations in Excel through automated formatting and metadata embedding. Among them, visual annotation includes **Orange Fill**: Applied to all residue positions within predicted domain boundaries (fields 17-18 in domtblout) which is presented in algorithm 3. In case of embedded metadata, dynamic cell comments are generated for each domain’s start position, containing:

- `Domain name:` *description* (from field 22+ in domtblout)
- `Accession:` *accession* (field 1)
- `Coords:` *start-end* (fields 17-18)
- Uses `openpyxl`’s `PatternFill` with exact RGB color matching
- Implemented coordinates(start-end positions) in the comment are not according to HMMER’s gap-free sequence numbering. They are calculated based on aligned protein sequences(contains ‘-’ in sequences)
- Skips invalid positions (start *>* end or negative values)

##### Algorithm 3: Domain Highlighting in Excel Worksheet

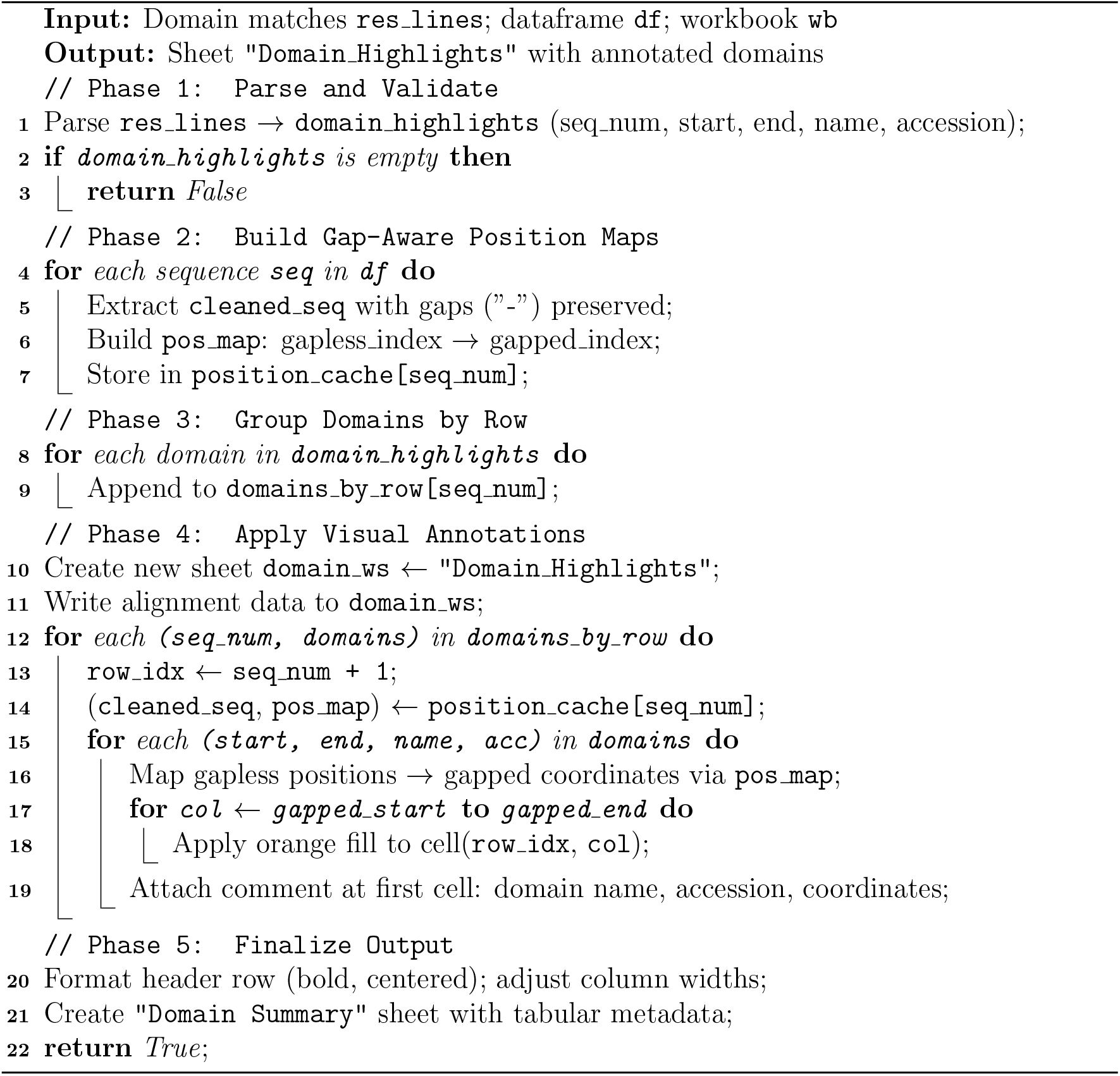

### 2.4 Motif–Domain Coverage Score (MDCS)

Motif detection and domain annotation are commonly performed as independent analyses. Yet sometimes position of some short linear motifs and conserved domain in protein structure or sequene provide us important biological insights. To support this interpretation, ProteoMapper computes the Motif–Domain Coverage Score (MDCS), a quantitative measure describing the spatial relationship between motifs and predicted protein domains.

For each motif occurrence and each overlapping Pfam domain within the same sequence, MDCS is defined as the proportion of the motif length that overlaps the domain:

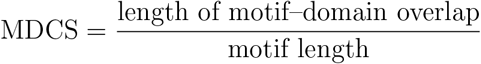

MDCS values range from 0 to 1. A value of 1 indicates that the motif is fully contained within a domain, values between 0 and 1 indicate partial overlap with domain boundaries, and a value of 0 indicates that the motif lies outside annotated domains. When a motif occurs multiple times within a sequence or overlaps multiple domains, ProteoMapper retains the highest MDCS value observed for each domain accession to summarize the strongest motif–domain association. MDCS calculation is performed using gapless sequence coordinates derived from `hmmscan` domain predictions to ensure consistency with Pfam annotations. The results are reported in a dedicated `MDCS Summary` worksheet in the output Excel file, listing the sequence index, motif pattern, associated domain accession(s), MDCS values, and a qualitative interpretation.

By summarizing motif–domain embedding in a numerical and easily filterable form, MDCS enables users to rapidly distinguish motifs that are likely part of core domain function from motifs that occur at domain boundaries or outside conserved domains. This information complements motif conservation and frequency analyses, providing an additional layer of context for downstream biological interpretation without adding substantial computational overhead.

### 2.5 Software and Tools Used

ProteoMapper integrates Python 3.10 with specialized libraries for protein analysis. Python packages such as, Pandas was used for data handling, NumPy was used for numerical operations, and openpyxl was used for Excel formatting. Domain detection employs HMMER v3.4 (hmmscan) with the Pfam-A.hmm v37.0 database, providing comprehensive domain annotation capabilities.

## 3 Results

This section discusses about different results from experiments, software interface, comparisons, and case studies.

### 3.1 User Interface

After running the project code, the interface shown in Figure 2 is displayed, providing an intuitive platform for users to input protein sequence alignments, configure parameters (e.g., regex patterns, conservation thresholds, and HMMER options), and visualize results. The interface allows real-time preview of motif matches, domain annotations, and user-defined highlights.

**Figure 2:**
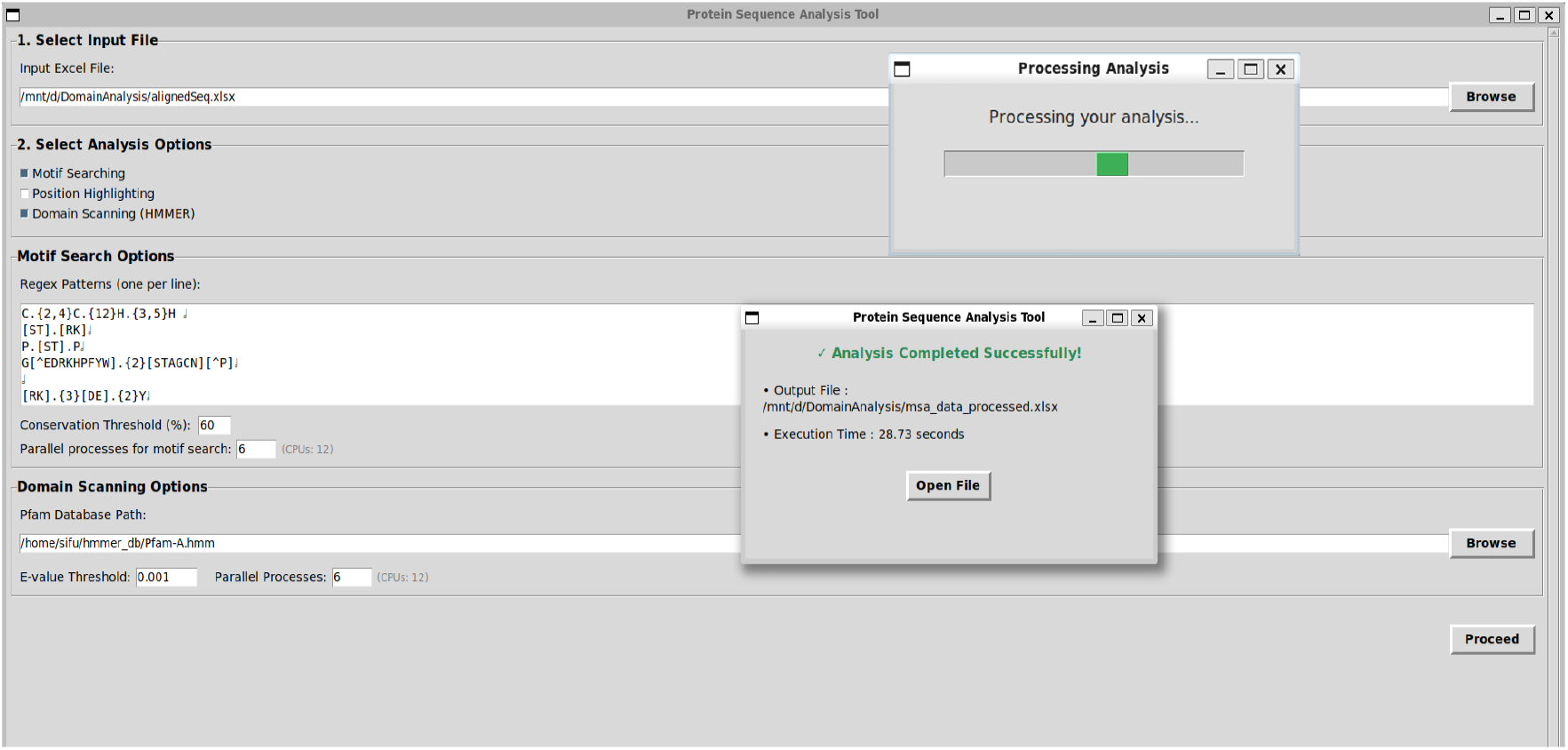
User interface of the Integrated Computational Toolkit, showing input fields, processing tab, and completion tab.

### 3.2 Dataset and Toolkit Output Structure

ProteoMapper was evaluated using three validated biological datasets: (i) 153 PLATZ transcription factor sequences from multiple plant species Qu et al. [25], Azim et al. [2], (ii) 44 actin-depolymerizing factor (ADF/cofilin) proteins Bamburg [4], Khatun et al. [12], and (iii) 17 ERD6-like (EDL) sugar transporter proteins from *Arabidopsis thaliana*, Kiyosue et al. [13], Slawinski et al. [27]. Each dataset was provided as a gapped multiple-sequence alignment in Excel format with associated gene name or gene ID columns. After processing, ProteoMapper produces a single multi-sheet Excel workbook that consolidates all analyses (Figure 3). The structure of the output is described below.

- **First Sheet:** Contains the complete alignment matrix. Regex-based motif matches are shown in **sky-blue**. Columns where a motif appears in ≥ 60% (threshold in this test) of sequences (user-adjustable) receive a **thick red border**, highlighting highly conserved motif positions without manual inspection (Figure 3D).
- **User-defined position highlighting:** Any alignment columns specified by the user (space-separated numbers) are filled bright green across all rows, enabling rapid visual mapping of known functional or variant sites (Figure 3D).
- **Match Summary:** An automatically generated table listing each user-defined regex pattern, its total match count, and all positional spans detected in the alignment. Column-wise frequencies facilitate identification of recurrent or conserved motif regions (Figure 3C).
- **Domain Highlights:** A duplicate of the alignment with Pfam domains obtained from parallel `hmmscan` scans (E-value ≤ 0.001 for this test). Detected domains are displayed in **orange**. Each domain block includes an embedded Excel comment containing the domain name, accession ID, bit score, E-value, and conditional E-value, providing complete HMMER metadata directly within the sheet (Figure 3B).
- **Domain Summary:** A tabular summary of all detected domains, including E-values, conditional E-values, and bit scores. E-values represent the expected number of false positives, with lower values indicating higher confidence, while bit scores provide a log-scaled measure of match quality (Figure 3E).
- **MDCS Summary:** A motif–domain relationship table reporting the Motif–Domain Coverage Score (MDCS) for each detected motif occurrence. For every sequence and motif pattern, the sheet lists the highest MDCS value observed for each overlapping Pfam domain (reported by accession ID), together with a qualitative interpretation (e.g., fully embedded or partial overlap). This summary enables rapid assessment of whether motifs are consistently embedded within conserved domain boundaries, as demonstrated in the ERD6-like protein case study (Figure 3A).
- **hitdata.txt:** A plain-text file containing all raw `hmmscan` hits, supporting downstream filtering, custom parsing, or archival of domain prediction results with complete metadata.

This unified Excel-based structure enables immediate comparison of motif occurrences, positional conservation, user-specified functional sites, conserved domain boundaries, and motif–domain embedding relationships. Analyses that previously required multiple independent tools and custom scripts are now integrated into a single, coherent output environment. Processed output files for all datasets are available at https://github.com/sifullah0/ProteoMapper/tree/main/sample_datasets%26outputs.

**Figure 3:**
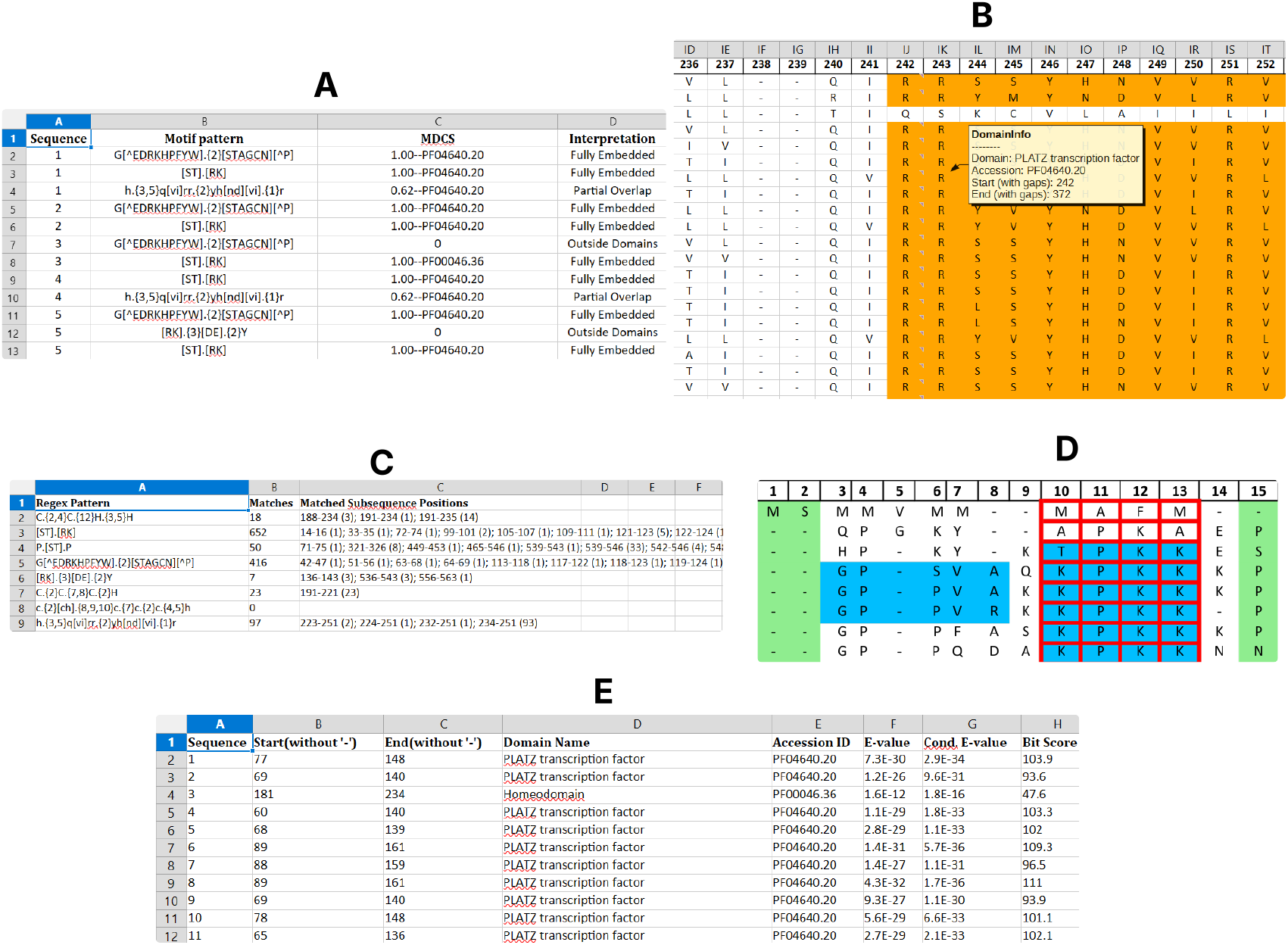
ProteoMapper output on representative datasets. (D) First Sheet showing skyblue motif matches and red borders on conserved columns; (C) Match Summary table; (B) Domain Highlights sheet with orange Pfam domains and hoverable metadata; (E) Domain Summary containing HMMER statistics; (A) MDCS Summary illustrating motif– domain embedding relationships.

### 3.3 Case Study-1: Validation of Domain Detection on BrPLATZ Proteins from *Brassica rapa*

To evaluate the accuracy of our protein analysis tool, we tested it using a published dataset of 24 PLATZ transcription factor proteins (BrPLATZ1–BrPLATZ24) from *Brassica rapa* L., as reported by Azim et al. [2]. The original study identified these proteins and analyzed their conserved domains using the SMART database [16]. The dataset included several domains such as PLATZ (PF04640), homeobox (PF00046), and B-box zinc finger (PF00643). The toolkit was applied to the same sequences using HMMER (`hmmscan`) with the Pfam-A database, using an E-value threshold of 0.001 and four CPU cores. The results were saved in the file `hitdata.txt` and presented in Table 1, which recorded all significant domain hits. ProteoMapper reproduced most of the reported domain annotations with high accuracy. For the PLATZ domain (PF04640), 22 of the 23 reported hits matched exactly, such as BrPLATZ1 (77–148) and BrPLATZ18 (133–204). One protein, BrPLATZ17, showed a shorter match (74–98 instead of 74–127). Neither analyses did not detected any PLATZ domain in BrPLATZ3. For the homeobox domain (PF00046), the tool detected it only in BrPLATZ3 (181–234), closely matching the reported range (180–235). No other proteins contained this domain.

**Table 1:** Comparison of PLATZ and Homeobox Domain Ranges Detected by SMART [2] and Our Tool (Pfam/HMMER).

| Protein | PLATZ<br>(Paper) | PLATZ<br>(Tool) | Homeobox<br>(Paper) | Homeobox<br>(Tool) | Match Type |
| --- | --- | --- | --- | --- | --- |
| BrPLATZ1 | 77–148 | 77–148 |  |  | Exact |
| BrPLATZ2 | 69–140 | 69–140 |  |  | Exact |
| BrPLATZ3 |  |  | 180–235 | 181–234 | Partial |
| BrPLATZ4 | 60–140 | 60–140 |  |  | Exact |
| BrPLATZ5 | 68–139 | 68–139 |  |  | Exact |
| BrPLATZ6 | 89–161 | 89–161 |  |  | Exact |
| BrPLATZ7 | 88–159 | 88–159 |  |  | Exact |
| BrPLATZ8 | 89–161 | 89–161 |  |  | Exact |
| BrPLATZ9 | 69–140 | 69–140 |  |  | Exact |
| BrPLATZ10 | 78–148 | 78–148 |  |  | Exact |
| BrPLATZ11 | 65–136 | 65–136 |  |  | Exact |
| BrPLATZ12 | 68–139 | 68–139 |  |  | Exact |
| BrPLATZ13 | 90–164 | 90–164 |  |  | Exact |
| BrPLATZ14 | 89–161 | 89–161 |  |  | Exact |
| BrPLATZ15 | 84–158 | 84–158 |  |  | Exact |
| BrPLATZ16 | 90–164 | 90–164 |  |  | Exact |
| BrPLATZ17 | 74–127 | 74–98 |  |  | Partial |
| BrPLATZ18 | 133–204 | 133–204 |  |  | Exact |
| BrPLATZ19 | 88–160 | 88–160 |  |  | Exact |
| BrPLATZ20 | 69–140 | 69–140 |  |  | Exact |
| BrPLATZ21 | 60–141 | 60–141 |  |  | Exact |
| BrPLATZ22 | 67–138 | 67–138 |  |  | Exact |
| BrPLATZ23 | 66–137 | 66–137 |  |  | Exact |
| BrPLATZ24 | 73–143 | 73–143 |  |  | Exact |

In addition to domain ranges, ProteoMapper provided detailed metrics such as bit scores (ranging from 28.3 to 111.0 for PLATZ hits) and E-values (all ≤ 7.3×10^−6^), confirming the statistical significance of the detections. The output file `hitdata.txt` also included conditional E-values and domain descriptions for further validation.

SMART reported possible B-box zinc finger domains (PF00643) in 15 of the 24 BrPLATZ proteins. However, the tool did not detect any significant B-box hits under the default E-value threshold (≤ 0.001). Pfam and SMART are both leading domain databases, but they are curated independently and use different hidden Markov models for the same domain, which can affect detection sensitivity [16]. Again, the SMART (Simple Modular Architecture Research Tool) tool identifies B-box domains with accessions such as SM00336 and Pfam (Protein Families Database) identifies B-box domains with accessions PF00643 (type 1) or PF07442 (type 2).

Pfam-based HMMER scan applies stricter thresholds based on well-curated alignments. Any hit with an E-value above 0.001 (for example, 0.005) is excluded automatically. Studies on PLATZ proteins also describe only the PLATZ domain and do not treat the B-box as a separate functional region [30]. Therefore, the absence of B-box detection in our results represents a more conservative annotation.

### 3.4 Case Study-2: Validation of the ADF-H Domain in Tomato *SlADF* Proteins

Khatun *et al*. [12] conducted a comprehensive genome-wide analysis identifying 11 actin depolymerizing factor (ADF) genes in tomato (*Solanum lycopersicum*). The study characterized these proteins based on their expression patterns across different organs and developmental stages. All 11 proteins were confirmed to harbor the ADF-H domain (also described as “Cofilin/tropomyosin-type actin-binding protein”, Pfam ID: PF00241) via SMART and InterPro tools, with domain spans detailed in Table 2 of their study.

**Table 2:** Comparison of ADF-H Domain Boundaries in 11 Tomato *SlADF* Proteins.

| Protein | Locus ID | Domain Span<br>(Paper) | Domain Span<br>(Tool) | IoU |
| --- | --- | --- | --- | --- |
| SlADF1 | Solyc01g094400 | 12–137 | 7–132 | 0.93 |
| SlADF2 | Solyc01g111380 | 16–143 | 12–142 | 0.94 |
| SlADF3 | Solyc03g025750 | 12–137 | 8–135 | 0.94 |
| SlADF4 | Solyc04g011370 | 12–139 | 7–137 | 0.93 |
| SlADF5 | Solyc06g005360 | 12–137 | 7–136 | 0.96 |
| SlADF6 | Solyc06g035980 | 12–137 | 7–135 | 0.94 |
| SlADF7 | Solyc09g010440 | 12–139 | 7–138 | 0.96 |
| SlADF8 | Solyc09g072590 | 18–145 | 14–145 | 0.96 |
| SlADF9 | Solyc09g090110 | 18–145 | 13–143 | 0.93 |
| SlADF10 | Solyc10g017550 | 12–139 | 7–138 | 0.96 |
| SlADF11 | Solyc10g084660 | 12–139 | 7–138 | 0.96 |

**Table 3:** Conservation analysis of PROSITE sugar transport signatures in a 17-sequence ERD6-like alignment using a 55% conservation threshold.

| Motif | Total matches | Span with frequency | Conserved? |
| --- | --- | --- | --- |
| PS00216 | 13 | 123–140 (6), 357–375 (7) | No |
| PS00217 | 10 | 165–190 (10) | Yes |

To validate the accuracy and reliability of this custom tool, the domain presence in *SlADF* protein sequences were reanalyzed. Sequences from the supplementary materials of Khatun *et al*. [12] were preprocessed (gap removal) and scanned via `hmmscan` against Pfam-A v35.0, with an E-value threshold of 0.001. Here, Domain scanning results for the first 11 sequences (the 11 *SlADFs*) were focused, comparing domain positions and overall concordance with Khatun *et al*. [12].

The ADF-H domain (PF00241.26) was detected in all 11 *SlADF* proteins with high confidence. Domain span overlap was quantified using **Intersection-over-Union (IoU)**, defined as:

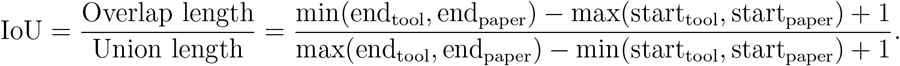

An IoU of 1 indicates perfect alignment; values *>* 0.9 indicate excellent overlap.

Domain boundaries showed strong concordance with the published spans, with a **mean IoU = 0.94** (range: 0.93–0.96), indicating *>* 90% positional overlap (Table 2). Start positions were slightly earlier in our tool (mean offset: −4.8 aa), likely due to Pfam’s focus on core conserved regions versus broader SMART annotations.

This analysis fully validates the annotations of the ADF-H domain in Khatun *et al*. [12], with 100% detection and near-perfect positional alignment (mean IoU 0.94). Minor boundary shifts reflect methodological differences (Pfam HMM vs. SMART) but do not affect functional interpretation. The tool provides a rapid, reproducible method for domain validation, suitable for large-scale plant genomic studies and integration with motif or expression analyses.

### 3.5 Case Study 3: Conservation and domain-context analysis of sugar transporter signatures in ERD6-like proteins

ERD6 was originally identified in *Arabidopsis thaliana* as an early dehydration-induced gene encoding a putative sugar transporter based on sequence similarity to known monosaccharide transporters [13]. Subsequent evolutionary analyses demonstrated that ERD6-like (ERDL/EDL) proteins form a conserved transporter family in land plants, while also exhibiting subfamily-specific sequence divergence [27]. Curated databases annotate ERD6-like proteins as members of the Major Facilitator Superfamily (MFS) and report the presence of two PROSITE sugar transport protein signatures, **PS00216** and **PS00217** [22, 23, 24, 28].

Seventeen ERD6-like/EDL proteins from *A. thaliana* (UniProt entries) were aligned and analyzed using ProteoMapper. Two PROSITE-derived motifs were searched using regular expressions corresponding to PS00216 and PS00217. Motif detection was performed on gapless sequences and projected back onto the gapped alignment for visualization. Domain annotation was carried out using HMMER against the Pfam database with a stringent E-value threshold of **1.0** *×* **10**^−5^. A conservation threshold of **55%** was applied, requiring a motif to occur at the same aligned position in at least 10 of 17 sequences to be classified as conserved.

Both sugar transporter signatures were detected. However, only PS00217 satisfied the conservation criterion. PS00217 was identified at a single aligned span (positions 165–190) in 10 of 17 sequences (58.8%) and was therefore classified as a conserved family-wide motif. In contrast, PS00216 produced a larger number of total matches (13), but these were distributed across two distinct aligned regions (positions 123–140 and 357–375), neither of which reached the conservation threshold. Consequently, PS00216 was not classified as conserved at 55%.

To further characterize the structural context of detected motifs, ProteoMapper computed the Motif–Domain Coverage Score (MDCS), which quantifies the fraction of a motif that overlaps a predicted Pfam domain. MDCS values close to 1 indicate fully embedded motifs within a domain, whereas values close to 0 indicate motifs located outside domain boundaries. Motifs with MDCS values close to 1 therefore suggest that they are likely integral to the core transport mechanism rather than peripheral or regulatory features. MDCS analysis showed that both PS00216 and PS00217 occurrences were predominantly fully embedded within predicted Pfam domain boundaries (MDCS=1). Despite both signatures are domain-embedded, only PS00217 as a family-wide conserved signature (58.8% frequency) and confirmed its role as an integral transport mechanism component, whereas PS00216 was identified as a non-conserved motif.

These results are consistent with the known biology of the ERD6-like family, which combines transporter features with lineage or subgroup-specific sequence variation [27]. Together, conservation analysis and MDCS-based domain context assessment demonstrate how ProteoMapper integrates motif detection, evolutionary conservation, and protein architectural interpretation within a unified workflow.

### 3.6 Performance Evaluation

ProteoMapper was benchmarked using 150 aligned PLATZ protein sequences and eight biologically relevant motif patterns: (i) `C.{2,4}C.{12}H.{3,5}H` (zinc finger C2H2), (ii) `[ST].[RK]` (PKC phosphorylation site), (iii) `P.[ST].P` (proline-directed phosphorylation), (iv) G[^EDRKHPFYW].{2}[STAGCN][^P] (N-myristoylation site), (v) `[RK].{3}[DE].{2}Y` (tyrosine kinase site), (vi) `C.{2}C.{7,8}C.{2}H` (B-box zinc finger), (vii) `C.{2}[CH].{8,10}C.{7}C.{2}C.{4,5}H` (NPC1-like cysteine-rich domain), and (viii) H.{3,5}Q[VI]RR.{2}YH[ND][VI].{1}R (PLATZ-specific motif). Two experiments were designed to assess scalability. First, how runtime grows with dataset size was measured by running the analysis on 50, 100, and 150 sequences using single process. Second, multiprocessing performance was evaluated by running the full 150-sequence dataset with 1, 2, 4, 6, and 8 processes.

The performance table demonstrates efficient scaling with dataset size and number of processes (Table 4). Runtime scales approximately linearly with sequence count in single-process mode, increasing from 10.9 s (50 sequences) to 29.0 s (150 sequences; Table 4). HMMER-based domain scanning dominates computational cost (∼93% of total runtime), whereas regex motif matching remains negligible (*<* 0.3 s).

**Table 4:** Performance scaling of ProteoMapper with sequence count and number of processes.

| Sequences | Processes | Motif (s) | Domain (s) | Total (s) | Speedup |
| --- | --- | --- | --- | --- | --- |
| 50 | 1 | 0.093 | 10.123 | 10.9 | – |
| 100 | 1 | 0.161 | 17.716 | 19.3 | – |
| 150 | 1 | 0.239 | 26.9 | 29.0 | 1.0× |
| 150 | 2 | 0.287 | 13.89 | 16.96 | 1.71× |
| 150 | 4 | 0.248 | 8.014 | 9.834 | 2.95× |
| 150 | 6 | 0.262 | 6.74 | 8.484 | 3.42× |
| 150 | 8 | 0.26 | 5.71 | 7.7 | 3.76× |

To quantify parallel efficiency, we define speedup as:

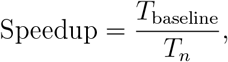

where *T*_baseline_ represents the runtime with one process and *T*_*n*_ represents the runtime with *n* processes. Parallel execution yields substantial improvements, reaching a maximum of 3.76*×* with eight processes. Parallel efficiency gradually declines beyond four processes due to increasing overhead from multiprocessing coordination, temporary file I/O, and the fact that each additional hmmscan instance loads the full Pfam database into memory. As a result, speedup rises sharply from 1*×* to ∼3*×* between 1 and 4 processes but plateaus thereafter, yielding only marginal gains (from 3.42*×* to 3.76*×*) when moving from 6 to 8 processes. Domain scanning benefits most from multiprocessing, improving almost fourfold from 26.9 s (single process) to 5.7 s (eight processes). In contrast, motif matching remains consistently low across all configurations, indicating minimal computational demand for pattern recognition tasks regardless of parallelization strategy.

## 4 Discussion

The results demonstrate that joint analysis of motif conservation and domain context provides biological insights that are difficult to obtain using domain- or motif-centric workflows alone. Across multiple protein families, ProteoMapper enabled systematic discrimination between motifs that are positionally conserved and structurally integrated within domains and those that exhibit positional variability despite sequence conservation. This distinction is biologically meaningful, as motifs that are both evolutionarily fixed and domain-embedded are more likely to contribute directly to core molecular mechanisms, whereas positionally flexible motifs may encode regulatory or lineage-specific functions. This principle is illustrated by the ERD6-like sugar transporter family. Two canonical PROSITE signatures, PS00216 and PS00217, were found to be fully embedded within conserved domains in all detected instances. However, only PS00217 exhibited strict positional conservation across the alignment, while PS00216 was distributed across multiple regions. The combination of equivalent domain embedding and differential positional conservation suggests functional subfunctionalization within a conserved structural scaffold. Such patterns are consistent with evolutionary models in which core transport mechanisms are constrained, while peripheral sequence features diversify to support regulatory or environmental adaptation. Importantly, this inference emerges directly from quantitative integration of conservation and domain context rather than from manual inspection or ad hoc comparison. Similar benefits were observed in PLATZ transcription factors and actin-depolymerizing factors, where conserved domain architectures were recovered with high accuracy and overlaid with motif conservation patterns. In these cases, integrated visualization facilitated rapid identification of conserved regulatory clusters and confirmation of architectural homogeneity across family members. Together, these examples demonstrate that integrated spatial analysis supports hypothesis generation related to regulatory mechanisms, evolutionary constraints, and functional divergence in protein families.

A central methodological contribution of this work is the introduction of quantitative metrics that formalize motif–domain relationships in an alignment-aware manner. Positional conservation scoring identifies motif occurrences fixed at identical alignment coordinates across a defined proportion of sequences, providing an explicit measure of evolutionary constraint on motif placement. Unlike frequency-based motif counts, this approach captures positional specificity, which is critical for distinguishing functionally constrained motifs from tolerated sequence patterns. The Motif–Domain Coverage Score (MDCS) further extends this framework by quantifying the fraction of a motif embedded within predicted Pfam domains. By expressing motif–domain overlap as a continuous value between 0 and 1, MDCS enables systematic classification of motifs as fully embedded, boundary-spanning, or extradomain. This abstraction transforms descriptive annotation into a quantitative variable that can be filtered, summarized, or correlated with other evolutionary or functional measures. Importantly, MDCS is not intended as a predictive score in isolation but as an interpretive metric that complements conservation analysis. Used together, positional conservation and MDCS distinguish motifs that are structurally constrained and evolutionarily fixed from those that are structurally permissive or evolutionarily flexible. This distinction is relevant for studies of regulatory logic, protein architecture, and functional divergence, and provides a basis for prioritizing motifs or regions for downstream experimental validation or variant effect assessment.

ProteoMapper is designed to complement, rather than replace, existing protein analysis tools. InterProScan remains the most comprehensive platform for large-scale domain annotation through integration of multiple signature databases, but requires command-line expertise, substantial computational resources, and produces complex output formats that are not optimized for interactive interpretation [1]. HMMER is the standard for profile-HMM–based domain detection, yet its text-based outputs require additional downstream processing to support integrative analyses [31]. A structured comparison of ProteoMapper with established motif and domain analysis tools is provided in Supplementary Table S1.

Motif-focused tools address different analytical needs. ScanProsite provides convenient web-based detection of curated PROSITE signatures but does not support domain annotation or large-scale batch analysis [7]. EMBOSS Fuzzpro enables command-line pattern matching but lacks alignment-level visualization and domain-context interpretation [26]. Probabilistic motif detection frameworks such as Wregex and its later versions offer improved sensitivity for weak or degenerate motifs through weighted regular expressions and position-specific scoring matrices [20, 21]. However, these approaches require curated training data and generate continuous scores that are less intuitive to interpret across aligned protein families, and they do not integrate domain annotation or structured visualization. Alignment-based motif detection tools such as MAFin provide conservation-aware motif identification and support multiprocessing at the command line [18]. Nevertheless, MAFin focuses primarily on nucleic acid motifs and does not provide an integrated graphical workflow for protein domain and motif analysis. Other systems emphasize downstream interpretation rather than spatial integration. CoSMoS ranks motif occurrences by evolutionary conservation but relies on precomputed alignments, lacks domain annotation, and offers limited organism coverage [17]. GOmotif associates motifs with Gene Ontology terms, providing functional enrichment at the protein level, but depends on web-based execution and does not support alignment visualization or motif–domain spatial analysis [5]. SCANMOT models combinations of motifs and their spacing to infer remote homology but lies outside the scope of integrated domain annotation and does not offer a modern desktop interface [6]. Within this landscape, ProteoMapper occupies a distinct niche by integrating motif detection, domain annotation, conservation analysis, and spatial quantification within an accessible, alignment-centric workflow. Its primary contribution lies in enabling interpretive analysis of motif–domain relationships rather than in maximizing detection sensitivity or database coverage.

ProteoMapper has several limitations. It does not currently support weighted motif scoring, PSSM-based models, or conservation-ranked motif discovery, which may limit sensitivity for highly degenerate or novel motifs. Explicit modeling of complex multi-motif architectures, such as spacing-dependent combinatorial signatures, is not yet implemented. Functional enrichment analysis of motif-containing proteins is also beyond the current scope. Additionally, very large-scale proteome analyses may be more efficiently handled by command-line pipelines such as InterProScan or HMMER in high-performance computing environments. Future development will focus on optional integration of weighted motif scoring while preserving interpretability, extension of multi-motif architecture modeling to capture co-evolutionary spacing constraints, and incorporation of functional enrichment analyses. These extensions aim to broaden analytical scope without compromising accessibility or alignment-aware visualization. Overall, ProteoMapper provides a practical and reproducible framework for integrated analysis of protein architecture, regulatory organization, and evolutionary constraint in functional genomics and comparative protein studies.

## 5 Conclusion

ProteoMapper introduces a quantitative framework for investigating the spatial and evolutionary relationships between conserved protein domains and short linear motifs. It utilizes the power of parallel processing in multiple cores for processing multiple sequences simultaneously to find large domains with a simple search for small motifs. Validation across 52 proteins from three families demonstrated both technical accuracy (mean domain boundary intersection-over-union of 0.94 and 100% detection of ADF-H domains) and biological insight generation. In ERD6-like sugar transporters, MDCS analysis revealed that canonical PROSITE signatures PS00216 and PS00217 are equally domain-embedded (MDCS = 1.0) yet evolutionarily distinct; PS00217 exhibits strict positional conservation (58.8% of sequences at positions 165–190), whereas PS00216 shows positional dispersion across two alignment regions. This pattern suggests functional subfunctionalization within a conserved structural framework. ProteoMapper complements specialized resources such as HMMER, ScanProsite, and the MEME Suite by providing an interpretive layer that links motif occurrence, domain context, and evolutionary constraint. By enabling systematic discrimination between domain-integrated and peripheral motifs, the framework supports investigation of regulatory mechanisms, evolutionary constraints, and functional divergence in protein families. These capabilities are particularly relevant for prioritizing sequence variants that disrupt conserved, domain-embedded motifs, which are more likely to affect core protein function than variants in unconstrained regions. The platform operates directly on Excel-formatted multiple-sequence alignments and produces interpretable, multi-sheet visual outputs without requiring programming expertise, supporting reproducible and iterative hypothesis testing. Future development will focus on integrating weighted motif scoring, extending multi-motif architecture modeling to capture co-evolutionary spacing constraints, and incorporating functional enrichment analyses. By integrating motif detection, domain annotation, conservation analysis, and quantitative spatial metrics within a unified workflow, ProteoMapper enables accessible investigation of protein architecture, regulatory organization, and evolutionary dynamics across functional genomics, comparative protein analysis, and structural bioinformatics.

## 6 Declaration of generative AI and AI-assisted technologies in the manuscript preparation process

During the preparation of this work the authors used ChatGPT and Claude in order to language refinement, grammar correction, and improving clarity in the manuscript. After using this tool/service, the authors reviewed and edited the content as needed and take full responsibility for the content of the published article.

## 7 Author Contributions

**Sifullah Mahmud:** Conceptualization; Methodology; Software; Formal analysis; Investigation; Data curation; Validation; Visualization; Writing – original draft. **Joyeeta Sarkar:** Data curation; Software (testing and debugging); Validation; Writing – original draft, review & editing. **Arif Hasan Khan Robin:** Conceptualization (biological motivation); Resources (datasets); Supervision; Writing – review & editing. **Machbah Uddin:** Conceptualization (project direction); Supervision; Writing – review & editing. All authors read and approved the final manuscript.

## 8 Acknowledgements

The authors acknowledge Bangladesh Agricultural University (BAU) for providing the academic and computational environment necessary to conduct this research. We are also grateful to the developers and maintainers of open-source resources, including HMMER and the Pfam database, which made this work possible.

